# Insulin resistance and adipose tissue inflammation induced by a high-fat diet are attenuated in the absence of hepcidin

**DOI:** 10.1101/2021.09.12.459942

**Authors:** Jithu Varghese James, Joe Varghese, Nikhitha Mariya John, Jean-Christophe Deschemin, Sophie Vaulont, Andrew T. McKie, Molly Jacob

## Abstract

Increased body iron stores and inflammation in adipose tissue have been implicated in the pathogenesis of insulin resistance (IR) and type 2 diabetes mellitus. However, the underlying basis of these associations are unclear. In order to assess this, we studied how IR and associated inflammation in adipose tissue developed in the presence of increased body iron stores. Male hepcidin knock-out (*Hamp1*^-/-^) mice, which have increased body iron stores, and wild-type (WT) mice were fed a high-fat diet (HFD) for 12 and 24 weeks. Development of IR and metabolic parameters linked to this, insulin signaling in tissue, and inflammation and iron-related parameters in visceral adipose tissue were studied in these animals. HFD-feeding resulted in impaired glucose tolerance in both genotypes of mice. In response to the HFD for 24 weeks, Hamp1-/- mice gained less body weight and developed less IR than corresponding WT mice. This was associated with less lipid accumulation in the liver and decreased inflammation and lipolysis in the adipose tissue in the knock-out mice, than in the WT animals. Fewer macrophages infiltrated the adipose tissue in the knockout mice than in wild-type mice, with these macrophages exhibiting a predominantly anti-inflammatory (M2-like) phenotype. These observations suggest a novel role of hepcidin (central regulator of systemic iron homeostasis) in the development of inflammation in adipose tissue and insulin resistance, in response to a high-fat diet.

**CLINICAL PERSPECTIVES:**

- Elevated body iron stores and inflammation in adipose tissue have been implicated in the pathogenesis of insulin resistance (IR) and type 2 diabetes mellitus. However, the underlying molecular mechanisms linking them are unclear.
- In response to high-fat diet (HFD)-feeding (to induce IR), mice that lacked hepcidin (*Hamp1*^*-/-*^) (and hence had elevated body iron stores) gained less body weight and developed less insulin resistance than wild-type (WT) mice. Inflammation and infiltration of macrophages into adipose tissue of HFD-fed *Hamp1*^*-/-*^ mice were less than in WT mice, with the macrophages exhibiting an anti-inflammatory M2-like phenotype.
- These findings suggest a novel role of iron and hepcidin in HFD-induced inflammation in adipose tissue and development of insulin resistance. They raise the possibility that modulation of body iron may represent a potential way to inhibit these processes.

## INTRODUCTION

Epidemiological studies have consistently shown a strong association between high serum ferritin levels (a marker of body iron stores) and increased risk of type 2 diabetes mellitus (T2DM) (1–4). Depletion of body iron stores (by phlebotomy or iron chelation) has been shown to decrease insulin resistance (IR), in patients with impaired glucose tolerance and T2DM (5,6). This suggests a causal association between increased body iron stores and IR. However, mechanisms underlying this association are poorly understood.

Inflammation in the visceral adipose tissue (VAT) has been postulated to be involved in the pathogenesis of obesity-induced IR (7,8). In mice fed a high-fat diet (HFD), inflammation is associated with infiltration of immune cells, especially macrophages, into the VAT, and secretion of pro-inflammatory cytokines (9). This is accompanied by a “phenotypic switch” in adipose tissue macrophages (ATMs), from an anti-inflammatory (M2-like) phenotype to a pro-inflammatory (M1-like) one (10). Sustained, low-grade inflammation induces lipolysis in VAT, resulting in ectopic lipid accumulation in the liver and skeletal muscle (11). These events have been postulated to play major roles in obesity-induced development of IR and T2DM (12,13)

Hepcidin is the chief regulator of systemic iron homeostasis (14). It binds to and degrades ferroportin, the cellular iron export protein, thus limiting intestinal iron absorption and macrophage iron recycling (15). Hereditary hemochromatosis (HH) comprises a group of conditions characterized by decreased hepcidin levels. Classical hemochromatosis, the most common form, is associated with iron overload and the classical triad of bronzed skin, cirrhosis and diabetes (16). The high incidence of diabetes in these patients has been attributed to an impairment in insulin secretion, rather than development of IR (17). In fact, patients with HH have been shown to exhibit increased insulin sensitivity (17). Similarly, mouse models of HH have shown supra-normal glucose tolerance and insulin sensitivity, despite severe iron accumulation in tissues (18). The mechanisms that underlie this are unclear.

Although hepcidin is primarily derived from the liver, adipose tissue has been reported to be an additional source, especially in obesity (19,20). Whether liver- or adipose tissue-derived hepcidin plays a role in the regulation of iron homeostasis in adipose tissue is not known. The iron content in macrophages has been shown to play a key role in determining macrophage polarization (21–23). However, it is not known whether hepcidin regulates iron content of resident and infiltrating macrophages in the adipose tissue. In this context, macrophages in patients with HH are iron-depleted (due to ferroportin overexpression) (24); whether this plays a role in increased insulin sensitivity observed in HH is not known.

In summary, elevations in body iron stores appear to be strongly associated with increased risk of developing IR. However, patients with HH have been reported to have enhanced insulin sensitivity. The mechanism that underlies these apparently contradictory findings and their interplay with inflammatory processes in adipose tissue is unclear. In order to study this, we used hepcidin knockout (*Hamp1*^-/-^) mice, a model of increased body iron load, to examine the effect of high-fat diet-induced effects on IR and inflammation in VAT.

## MATERIALS AND METHODS

### Experimental animals

The hepcidin knockout (*Hamp1*^*-/-*^) mice were global knock-out mice and the generation of these mice has been previously reported (25). Male hepcidin knockout mice (*Hamp1*^-/-^ mice) (C57Bl/6J), which were iron-overloaded (25), were used for the study; wild type (WT) mice served as control animals. All experiments were carried out in accordance with the regulations of the Committee for the Purpose of Control and Supervision of Experiments on Animals (CPCSEA), Government of India.

At 7 weeks of age, mice of both genotypes were shifted from regular chow diet (Diet no. #D131, SAFE diets, France) to a control diet (CD) (#D12450J, 10% of total calories from fat; Research Diets, Inc., USA). One week later (at 8 weeks of age), the mice were randomly allocated to receive a high-fat diet (HFD) (#D12492, 60% of total calories from fat; Research Diets Inc., USA) or continue on the CD for 12 or 24 weeks, with access to food and water *ad libitum*. The iron content of the CD and HFD were 43 mg Fe/ kg and 58 mg Fe/kg respectively.

Mice were housed in standard cages (290 × 220 × 140 mm), with each cage housing two animals. Suitable environmental enrichment was provided in these cages. Mice were maintained under appropriate temperature and humidity control. Calorie intake and weight gain were monitored weekly. After the completion of the periods of feeding (12 or 24 weeks), mice were euthanized under terminal anesthesia (intra-peritoneal injection of xylazine and ketamine at a dose of 100 mg/kg and 10 mg/kg body weight, respectively). Blood and tissue (the liver, epididymal white adipose tissue [eWAT] (representative of visceral adipose tissue in rodents (26)), inguinal pad of subcutaneous adipose tissue [iSAT], spleen and quadriceps muscle) were harvested; snap-frozen in liquid nitrogen and stored at -70°C, until further processing. Euthanasia of the mice were carried out between 08 to 11 hours on the days concerned.

Insulin signaling was studied in a subset of both genotypes of mice fed CD or HFD for 24 weeks. For this, at the end of 24 weeks, mice were fasted for 10 hours overnight and then injected with insulin (Actrapid, Novo Nordisk), intraperitoneally (1U/kg) (27). Seven minutes later, they were euthanized. Samples of blood, liver, eWAT and quadriceps muscle were harvested. In order to study the baseline characteristics, 8-week-old WT and *Hamp1*^-/-^ mice (n=3-4) were used.

### Glucose and insulin tolerance tests

Glucose tolerance tests (GTT) and insulin tolerance tests (ITT) were carried out in the mice, one week prior to euthanasia respectively, using standard protocols (28). For the GTT, mice were fasted for 6 hours before the test. Glucose was administered at a dose of 2g/kg body weight, by intra-peritoneal injection. A blood sample was obtained from the tail vein of each mouse, before administering the glucose (0 mins). After administration of glucose, blood samples were collected from each mouse, at 15, 30, 60 and 120 min after the glucose load. Glucose levels in these samples were determined, using a glucometer (Bayer Contour, USA). For the ITT, mice were fasted for 4 hours; insulin (Actrapid insulin, Novo Nordisk, Denmark) was injected intra-peritoneally at a dose of 0.75 U/kg body weight. Glucose levels were estimated in blood samples obtained from the tail vein immediately before the insulin injection (at 0 mins) and at 15, 30, 45 and 60 min after the insulin injection. For the GTT and ITT, the blood glucose values obtained for each mouse was plotted against time. The area under curve (AUC) was determined, using the linear trapezoid method.

### Estimation of serum glucose and insulin levels

Glucose (GOD/POD method) and insulin (Mercodia, Sweden) were estimated in serum. The homeostatic model assessment (HOMA)-IR index was calculated using the following formula:

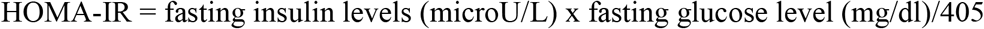

### Estimation of cytokines

Levels of tumor necrosis factor alpha (TNFα) and interleukin-6 (IL-6) were estimated in eWAT lysates and in serum by ELISA (TNFα-DY410 and IL-6 DY406, R & D Systems, Minneapolis, USA).

### Flow cytometric analyses

Adipose tissue macrophages (ATMs) were isolated and characterized as described previously, with certain modifications (29). Briefly, the stromal vascular fraction (SVF) was isolated from eWAT by collagenase digestion. The cells were then incubated with CD16/CD32 blocking reagent (Miltenyi Biotech, Germany) (4°C for 5 min), and with fluorescence-labeled primary antibodies (4°C for 30 min). Antibodies used were anti-F4/80-FITC, anti-CD11b-Vioblue, anti-CD301-PE and CD11c-APC (Miltenyi Biotech, Germany) (Supplementary Table 4). FACS Aria III flow cytometer (BD Biosciences, USA) was used for data acquisition. Unstained, single-stained and fluorescence minus one (FMO) controls were used for compensation and data analyses. Representative images for the gating strategy employed are shown in the Supplementary figure 2. Events were first gated to identify singlet cells and exclude dead cells and cellular debris. Selected events were then gated to identify ATMs (cells that co-express F4/80 and CD11b [F4/80^+^ CD11b^+^]). F4/80^+^ CD11b^+^ cells (macrophages) that expressed CD301 (CD301^+^ CD11c^-^) were identified as anti-inflammatory (M2-like) macrophages and those that expressed CD11c (CD11c^+^ CD301^-^) were identified as pro-inflammatory (M1-like) macrophages. Data obtained were analysed using FlowJo, version 10.4 (FlowJo, LLC, USA).

### Lipolysis assay

Lipolysis in eWAT was determined *ex vivo* (30). Briefly, the fat pad was incubated in DMEM containing 1% BSA (free of fatty acids) and 5 μM triacsin C, at 37°C for 60 min, in a CO_2_ incubator. Free fatty acids released from the tissue was measured in the medium (WAKO, Japan) and was normalized to the protein content of the tissue explant.

### Tissue triglyceride estimation

Lipids in the liver and quadriceps muscle were extracted by Folch’s extraction procedure (31). Briefly, 100 mg of tissue were homogenized in 900 μL of PBS. The homogenate (200 μL) was then added to a mixture of chloroform and methanol (2:1 v:v) (1.2 mL) and mixed thoroughly. To this, 100 μL of PBS was added to make a final volumetric ratio of 8:4:3 (v/v/v) of chloroform/methanol/water. This was mixed vigorously and then centrifuged at 3000 rpm at 4°C for 10 min. The lower organic phase (200 μL), which separated out, was removed into a fresh tube and allowed to air-dry. To this tube, 1% Triton X-100 in ethanol was added. The dissolved lipid extract obtained at the end of this step was used to measure triglycerides, using a commercially available kit (Randox, UK).

### Tissue iron estimation

Iron content was estimated in the liver, skeletal muscle, spleen, eWAT and adipocytes (isolated from the eWAT) (32), by atomic absorption spectrophotometry (Perkin-Elmer AA200 atomic absorption spectrometer) (for eWAT and adipocytes) or by a bathophenanthrolene-dye binding method (for the rest). The iron content of the eWAT was also assessed by *in-situ* perfusion with Perls’ Prussian blue solution (22).

### Histological examination of liver and adipose tissue

Histological examination of the liver and eWAT were carried out, using hematoxylin and eosin and Perls’ staining for iron. The size and numbers of adipocytes in the eWAT were determined (33). In each section of adipose tissue examined, 20 high-power fields were studied, and 100 adipocytes were chosen in each of these fields. The size of these adipocytes was determined by measuring their diameters and areas, using the ImageJ (NIH, USA).

### Western blot analyses

Samples of liver, eWAT and quadriceps muscle were homogenized in RIPA buffer containing a cocktail of protease inhibitors. The lysates were centrifuged at 14,000*g* for 20 minutes to remove cellular debris. Protein content of the supernatants was estimated using the BCA method (Pierce, USA). These were then separated by SDS-PAGE (50 µg of protein) and transferred to polyvinylidene difluoride membranes. The membranes were blocked with 5% BSA (in TBS-T) and probed with appropriate primary antibodies overnight at 4°C. Following this, membranes were washed and incubated with appropriate secondary antibodies. Bands were detected using an enhanced chemiluminescence kit (SuperSignal West Dura, Thermo Fisher Scientific, USA) and quantified using ImageJ (NIH, USA). Beta (β)-actin was used as the loading control. Sources and dilutions of primary and secondary antibodies used are given in Supplementary Table 5.

### Quantitative real-time PCR

Tissue samples of the liver, eWAT and adipocytes were homogenized in Tri-Reagent and total RNA was isolated, according to the manufacturer’s instructions (Merck, Germany). One microgram of isolated RNA was used to synthesize cDNA, using the Reverse Transcriptase Kit (Takara, Japan). Quantitative PCR reactions were carried out in duplicate, using the SYBR Premix Ex Taq II (Takara, Japan), using a BioRad Chromo4 real-time PCR machine. The relative mRNA levels for the genes of interest were calculated using the 2^-ΔΔCt^ method, and normalized to *Hprt*, which was used as the reference gene. The list of primers used, and the primer validation data are given in Supplementary Table 6 and 7.

### Statistical analysis

Statistical Package for Social Scientists (SPSS), version 16.0, was used for data analyses. Normally distributed data were analysed using one-way ANOVA, followed by Tukey multiple comparison post-hoc test. Data with skewed distribution were analysed by the Kruskal–Wallis test, followed by Mann–Whitney U test for pair-wise comparisons. Correlational analysis was done by Spearman’s correlation test. A P value <0.05 was considered significant in all cases.

## RESULTS

Data from both wild-type and *Hamp1*^-/-^ mice at baseline (0 weeks of feeding) are shown in Supplementary Table 1.

### *Hamp1*^-/-^ mice gained less weight than WT mice, in response to HFD-feeding

Gains in body weight were similar in the two genotypes on the CD, after 12 and 24 weeks of feeding (Fig 1A to C). Increases in body weight in HFD-fed mice were higher than those in mice of the corresponding genotype on the CD. After 12 weeks of HFD feeding, the weight gains in WT and *Hamp1*^*-/-*^ mice were similar (Fig 1A and C). In the groups fed the HFD for 24 weeks, weight gains in the 2 genotypes were similar till about the 17^th^ week of feeding; subsequently, weight gains in the *Hamp1*^*-/-*^ mice were significantly lower than those in the WT mice (Fig 1B). After 24 weeks of HFD, the mean gain in body weight was significantly lower in *Hamp1*^*-/-*^ mice than in WT mice (Fig 1C). The average calorie intake by mice in the HFD-fed group was higher than that in the CD-fed mice, in both genotypes. Within the HFD-fed mice, calorie intake was similar in both genotypes, for both 12 and 24 weeks of feeding (Fig 1D).

**Figure 1:**
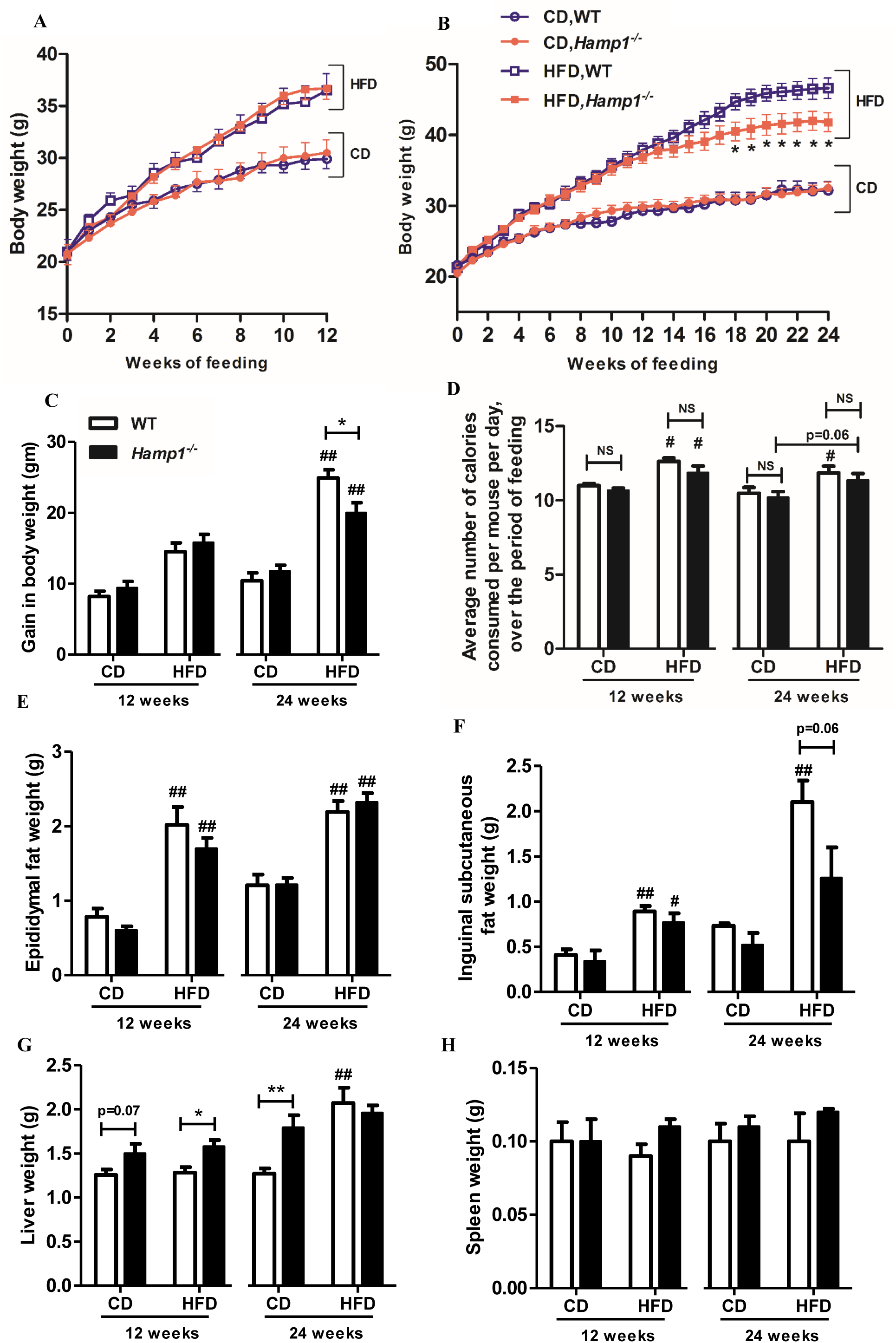
Effect of HFD feeding on body and tissue weights in *Hamp1*^-/-^ and WT mice. Body weights over 12 (A) and 24 (B) weeks of feeding, gains in body weight (C) and food consumption (D) of wild-type (WT) and *Hamp1*^*-/-*^ mice fed a control diet (CD) or high-fat diet (HFD), for 12 weeks or 24 weeks. Weights of epididymal white adipose tissue (eWAT) (E), inguinal subcutaneous adipose tissue (iSAT) (F), the liver (G) and spleen (H) were measured in these mice after the feeding period of 12 and 24 weeks. For (F) and (H) n = 6 mice; for all other variables, n = 10 to 12 mice. Results are shown as mean ± SE. For (C), (E) and (G) data were analysed by one-way ANOVA, followed by Tukey post-hoc test; for all the other variables Kruskal–Wallis test was used, followed by Mann–Whitney U test for pair-wise comparisons. \**P* < 0.05, *\*\*P* < 0.01 when compared with corresponding WT group; #*P* < 0.05, ##*P* < 0.01 when compared with corresponding genotype in the CD-fed group.

Weights of eWAT and iSAT in WT and *Hamp1*^*-/-*^ mice, fed the HFD, were higher than those from corresponding CD-fed mice, both at 12 and 24 weeks. While the weights of eWAT were similar in both genotypes after HFD feeding for 24 weeks (Fig 1E), weights of iSAT in *Hamp1*^*-/-*^ mice tended to be lower than in HFD-fed WT mice (p=0.06) (Fig 1F). After 12 weeks of feeding, liver weights in *Hamp1*^*-/-*^ mice were higher than in WT mice, irrespective of the diet fed. After 24 weeks of CD feeding, liver weights in *Hamp1*^*-/-*^ mice were significantly higher than in WT mice. HFD-feeding significantly increased the liver weight in WT mice, but this was not seen in *Hamp1*^*-/-*^ mice (Fig 1G). After 12 and 24 weeks of feeding, spleen weights were similar in WT and *Hamp1*^*-/-*^ mice, irrespective of the diet fed (Fig 1H).

### *Hamp1*^*-/-*^ mice developed impaired glucose tolerance similar to WT mice, in response to HFD-feeding, but less insulin resistance

On the CD, both WT and *Hamp1*^*-/-*^ mice showed similar responses for the Glucose tolerance tests (GTT) and insulin tolerance tests (ITT), after 12 and 24 weeks of feeding (Fig 2A to F). On HFD-feeding, both genotypes developed impaired glucose tolerance to similar extents (Fig 2A to C). After HFD-feeding, WT mice showed greater insulin resistance (AUC in the ITT), after 12 and 24 weeks of feeding, than CD-fed mice (Fig 2D to F). However, this was not seen in HFD-fed *Hamp1*^*-/-*^ mice, when compared with CD-fed *Hamp1*^*-/-*^ mice. In fact, at 24 weeks, the AUC for the ITT in the HFD-fed *Hamp1*^*-/-*^ mice was significantly lower than that of the WT mice on the HFD (Fig 2F). Thus, *Hamp1*^*-/-*^ mice developed less insulin resistance than WT mice, in response to HFD-feeding.Fasting serum insulin concentrations and the HOMA-IR index (a surrogate marker of insulin resistance), were significantly lower in HFD-fed *Hamp1*^*-/-*^ mice, compared to WT mice fed HFD for 24 weeks (Fig 2G and H). Fasting glucose levels were similar in both genotypes, irrespective of the diet they were on (Fig 2I).

**Figure 2:**
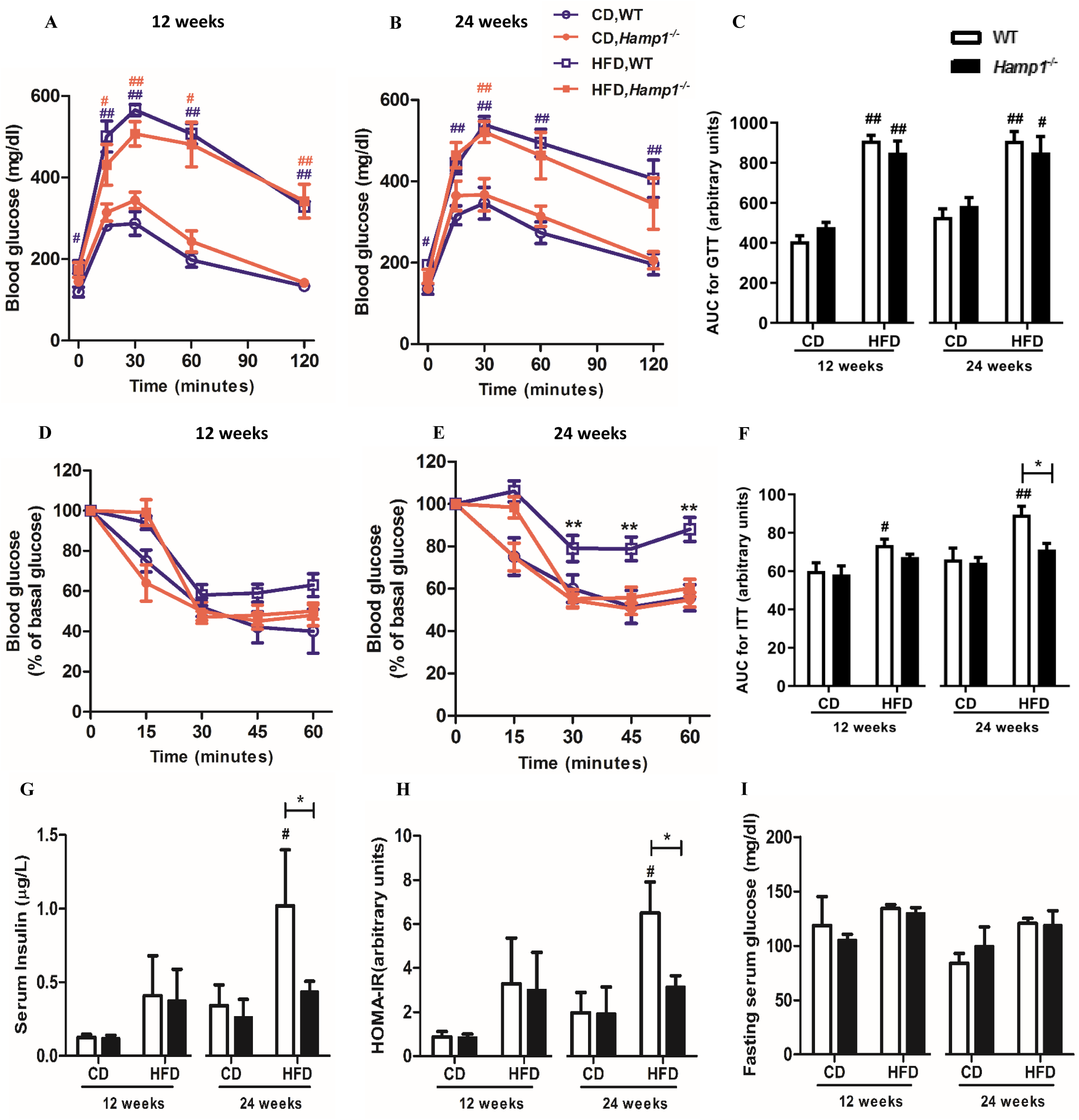
*Hamp1*^*-/-*^ mice developed impaired glucose tolerance, but less insulin resistance than WT mice, in response to 24 weeks of HFD-feeding. Intraperitoneal glucose tolerance test (GTT) (A and B) and insulin tolerance tests (ITT) (D and E) were done in WT and *Hamp1*^-/-^ mice fed a control diet (CD) or a high-fat diet (HFD) for 12 weeks (n = 6 in each group) (A and D) and 24 weeks (n = 6 for GTT and n = 12 for ITT) (B and E). The areas under the curves (AUC) for GTT (C) and ITT (F) were calculated by the linear trapezoid method. Serum insulin (G) and serum glucose (H) in WT and *Hamp1*^-/-^ mice fed a CD or HFD for 12 weeks and 24 weeks (n = 4 to 5). For (G) and (I) data were analysed by one-way ANOVA, followed by Tukey post-hoc test; for all the other variables Kruskal–Wallis test was used, followed by Mann–Whitney U test for pair-wise comparisons. Results are shown as mean ± SE; \**P* < 0.05 when compared with corresponding WT group; #*P* < 0.05, ##*P* < 0.01 when compared with corresponding genotype in the CD-fed group (blue=WT mice, red = *Hamp1*^-/-^ mice).

### HFD-induced macrophage infiltration and inflammation in eWAT were attenuated in *Hamp1*^*-/-*^ mice

After 12 weeks of feeding, gene expression of inflammatory cytokines and macrophage markers in the eWAT (representative of visceral adipose tissue in rodents (26)) were similar in both genotypes, irrespective of the diet fed (Supplementary Fig 1). After a HFD for 24 weeks, mRNA expression of inflammatory markers in the eWAT was higher in both genotypes than in corresponding CD-fed mice (Fig 3A and C). However, in the *Hamp1*^*-/-*^ mice, HFD-induced increases in mRNA levels of TNFα were significantly lower than that in the WT mice (Fig 3A). Levels of TNFα in the eWAT lysate from these mice also tended to be lower than in the corresponding WT mice (p = 0.06) (Fig 3B). IL-6 mRNA expression in HFD-fed *Hamp1*^*-/-*^ mice was significantly higher than in the HFD-fed WT mice, but its levels in tissue lysates were similar (Fig 3C and D).

**Figure 3:**
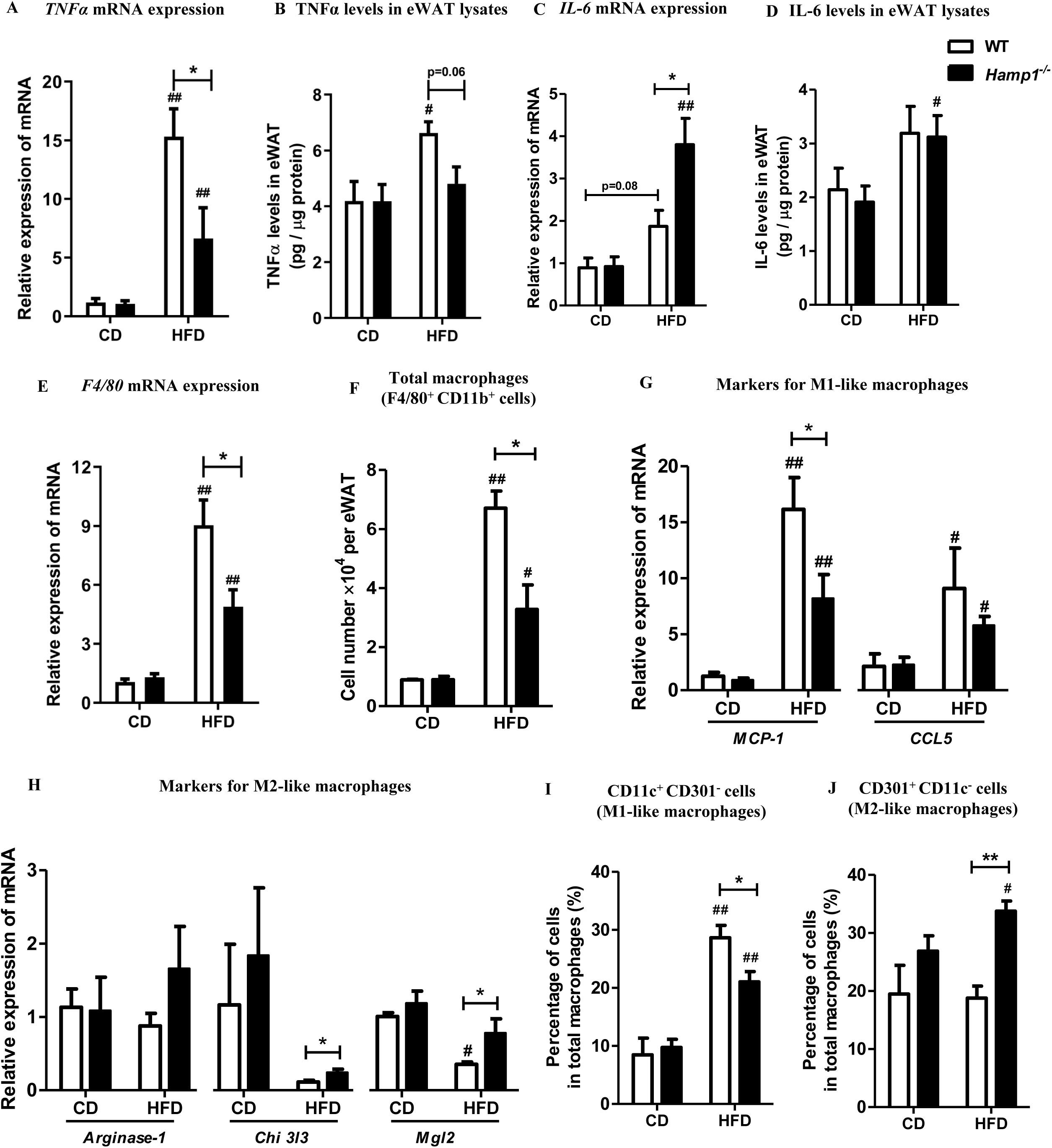
Inflammation in adipose tissue was attenuated in *Hamp1*^*-/-*^ mice fed HFD for 24 weeks. Expression of mRNA for tumor necrosis factor alpha (TNFα) (A) and interleukin-6 (IL-6) (C) in the eWAT of WT and *Hamp1*^-/-^ mice, fed a CD or HFD for 24 weeks. Protein expression of TNFα (B) and IL-6 (D) in the eWAT lysates from these mice. The mRNA levels of F4/80 (E), *MCP-1* and *CCL5* (markers of pro-inflammatory macrophages or M1-like macrophages) (G), *arginase 1, Chi3l3* and *Mgl2* (markers of anti-inflammatory macrophages or M2-like macrophages) (H) in the eWAT of WT and *Hamp1*^-/-^ mice fed a CD or HFD for 24 weeks. Relative expression was calculated by normalising expression of the gene of interest to that of the housekeeping gene, *Hprt*. F: Quantification of macrophages (F4/80+ CD11b+) in the eWAT of WT and *Hamp1*^-/-^ mice fed a CD or HFD for 24 weeks, as determined by flow cytometry. Quantification of pro-inflammatory (M1-like macrophages) (CD11c+ CD301-cells) (I) and anti-inflammatory (M2-like macrophages) (CD11c-CD301+ cells) (J) among the total macrophages in the stromal vascular fraction (SVF) from the eWAT of these mice. Results are shown as mean ± SE (n=5 to 6 in each group). Data were analyzed by Kruskal–Wallis test, followed by Mann–Whitney U test for pair-wise comparisons. \**P* < 0.05, *\*\*P* < 0.01 when compared with corresponding WT group; #*P* < 0.05, ##*P* < 0.01 when compared with corresponding genotype in the CD-fed group.

After 24 weeks of CD feeding, mRNA levels of *F4/80* (a macrophage marker), *Mcp-1* and *CCL5* (markers of pro-inflammatory [M1-like] macrophages) and *Arg1, Chi3l3* and *Mgl2* (markers of anti-inflammatory [M2-like] macrophages) were similar in both genotypes (Fig 3E, G and H). In response to HFD, *F4/80* expression was significantly higher in both the genotypes, when compared with their CD-fed counterparts, showing increased macrophage content in the eWAT. However, F4/80 mRNA level was significantly lower in HFD-fed *Hamp1*^*-/-*^ mice than in WT mice on the same diet (Fig 3E). At the same time, in HFD-fed *Hamp1*^*-/-*^ mice, mRNA levels of *Mcp-1* were significantly lower and those of *Chi3l3* and *Mgl2* were significantly higher in eWAT, than in corresponding WT mice (Fig 3G and H).

Consistent with the above results, flow cytometry analyses of the stromal vascular fraction (SVF) from eWAT showed that the number of macrophages (F4/80^+^ CD11b^+^ cells) were significantly lower in HFD-fed *Hamp1*^*-/-*^ mice than in WT mice on the same diet (Fig 3F). In addition, HFD-fed *Hamp1*^*-/-*^ mice had significantly fewer M1-like macrophages (CD11c+ CD301^-^ cells) and significantly more M2-like macrophages (CD11c-CD301+ cells) in the SVF fraction, than HFD-fed WT mice (Fig 3I and J). These results show that macrophage infiltration into eWAT was significantly lower, and that they had a predominantly anti-inflammatory phenotype, in *Hamp1*^*-/-*^ mice when compared to WT mice after 24 weeks of HFD-feeding. We also found that the mRNA expression of *Arg1* tended to be higher in the SVF from the *Hamp1*^*-/-*^ mice on HFD than WT on the same diet (p=0.08); mRNA expression of MCP-1 was similar in the 2 groups (Supplementary Fig 1I). Determination of additional markers was not possible because of the limited quantity of SVF obtained. Correlation analysis showed that insulin resistance (AUC for ITT) in HFD-fed WT mice positively correlated with mRNA levels of *CCL5* and tended to correlate with that of F4/80 (p=0.07) in eWAT, while these did not correlate in HFD-fed *Hamp1*^*-/-*^ mice (Supplementary Table 2).

### HFD-induced lipolysis in adipose tissue and hepatic steatosis were attenuated in *Hamp1*^*-/-*^ mice

After HFD-feeding for 24 weeks, *Hamp1*^*-/-*^ mice were found to have a higher number of smaller-sized adipocytes and fewer larger-sized adipocytes, than the HFD-fed WT mice (Fig 4A). However, values for the mean area of adipocytes were similar in both genotypes in the HFD-fed group (Fig 4B). Adipocyte sizes were similar in WT and *Hamp1*^*-/-*^ mice fed the CD (Supplementary Fig 3A and Fig 4B). Serum levels of free fatty acids (FFA) were also similar in both genotypes, irrespective of the type of diet fed (Fig 4C). However, the amount of FFA released *ex vivo* from the eWAT of HFD-fed *Hamp1*^*-/-*^ mice tended to be lower than that from WT mice, suggesting lower rates of lipolysis in the eWAT from *Hamp1*^*-/-*^ mice (p = 0.07) (Fig 4D).

**Figure 4:**
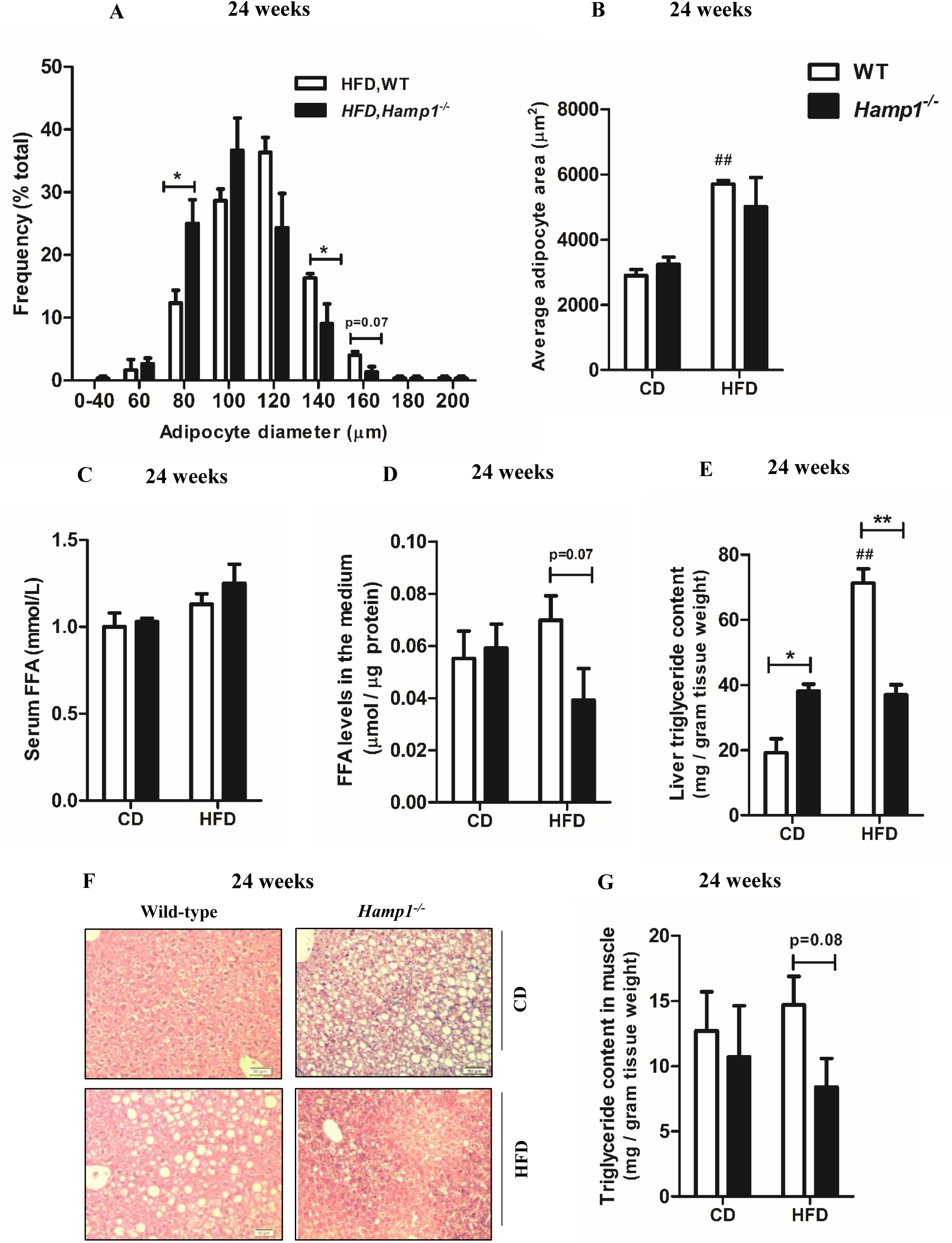
Changes in visceral adipose tissue, and ectopic lipid accumulation in the liver and skeletal muscle from WT and *Hamp1*^*-/-*^ mice, in response to 24 weeks of HFD-feeding. (A) Frequency distribution of adipocytes, based on their diameters, in WT and *Hamp1*^-/-^ mice fed a HFD for 24 weeks, n = 3 in each group. (B) Average area of adipocytes in the eWAT of WT and *Hamp1*^-/-^ mice fed a CD or HFD for 24 weeks, n = 3 in each group. (C) Serum levels of FFA in WT and *Hamp1*^-/-^ mice fed a CD or HFD for 24 weeks, n = 6 in each group. (D) Free fatty acid (FFA) concentrations in the incubation medium of eWAT (*ex-vivo*) from these mice were measured as an indicator of lipolytic activity, n = 3 in each group. Triglyceride content in the liver (E) and quadriceps muscle (G) of WT and *Hamp1*^-/-^ mice fed a CD or HFD for 24 weeks, n = 6 in each group. (F) Representative images of histological sections of the liver sections stained with hematoxylin and eosin and Perls’ staining (20X). Results are shown as mean ± SE. For (E) and (G) data were analysed by one-way ANOVA, followed by Tukey post-hoc test; for all the other variables Kruskal–Wallis test was used, followed by Mann–Whitney U test for pair-wise comparisons. \**P* < 0.05, *\*\*P* < 0.01 when compared with corresponding WT group; ##*P* < 0.01 when compared with corresponding genotype in the CD-fed group.

After 12 and 24 weeks of HFD-feeding, the hepatic triglyceride content in WT mice was significantly higher than in CD-fed WT mice (Fig 4E, Supplementary Fig 3B). *Hamp1*^*-/-*^ mice on the CD were found to have significantly higher levels of hepatic triglycerides than the WT mice on the same diet, after 12 and 24 weeks of feeding (Fig 4E, Supplementary Fig 3B). However, after HFD-feeding for 24 weeks (but not at 12 weeks), *Hamp1*^*-/-*^ mice had significantly lower hepatic triglyceride levels than corresponding WT mice (Fig 4E). These results were corroborated by findings of greater extent of hepatic steatosis in the CD-fed *Hamp1*^*-/-*^ mice than in CD-fed WT mice, and less so in HFD-fed *Hamp1*^*-/-*^ mice than in HFD-fed WT mice (Fig 4F). The triglyceride content in the quadriceps muscle also tended to be lower in *Hamp1*^*-/-*^ mice than WT mice, after 24 weeks of HFD feeding (p = 0.08) (Fig 4G).

### Insulin signaling in tissues from WT and *Hamp1*^*-/-*^ mice, after 24 weeks of HFD-feeding

In the liver of CD-fed *Hamp1*^*-/-*^ mice, levels of pAkt (Ser473) were significantly higher (both in the basal state [without insulin stimulation] and in response to insulin) than in the WT mice on the same diet (Fig 5A). In the eWAT and quadriceps muscle, basal and insulin-stimulated pAkt levels in the CD-fed mice were similar in both genotypes (Fig 5B and C). In mice fed the HFD, basal levels of pAkt were significantly higher only in the eWAT of *Hamp1*^*-/-*^ mice, compared to WT mice (Fig 5B). Insulin-stimulated increases in pAkt levels were similar is all 3 tissues from HFD-fed mice of both genotypes (Fig 5A to C).

**Figure 5:**
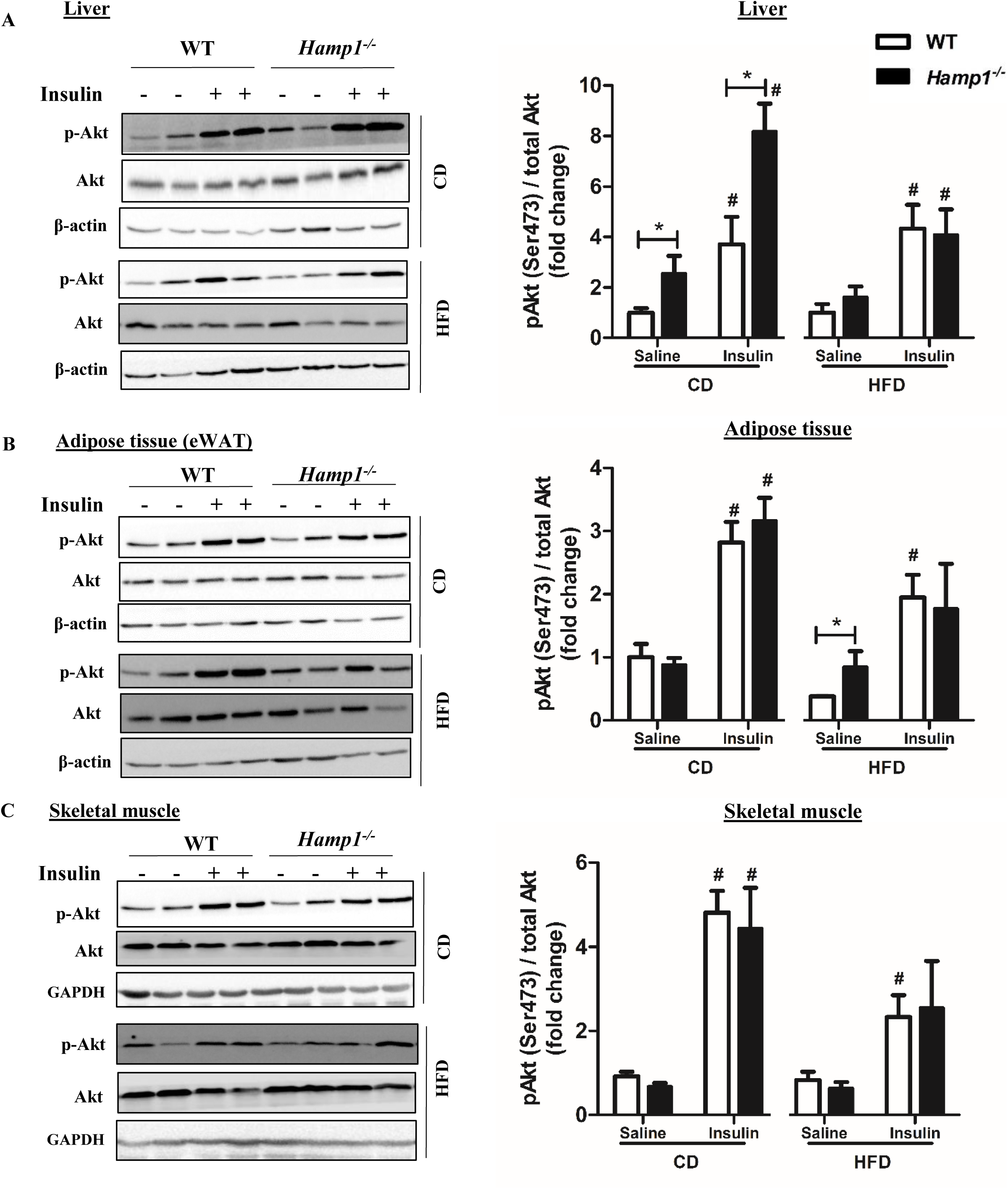
Insulin signaling in liver, adipose tissue and skeletal muscle in WT and *Hamp1*^*-/-*^ mice, in response to 24 weeks of HFD-feeding. Representative images and densitometric quantification of western blots for phosphorylated p-Akt, normalized to total Akt, in the liver (A), eWAT (B) and skeletal muscle (C) of WT and *Hamp1*^-/-^ mice, fed a CD or HFD for 24 weeks; n = 3 to 4 in each group. β-actin (for liver and eWAT) and GAPDH (for skeletal muscle) were used as loading controls. Results are shown as mean ± SE. Data were analysed by Kruskal–Wallis test, followed by Mann–Whitney U test for pair-wise comparisons. \**P* < 0.05 when compared with corresponding WT group; #*P* < 0.05 when compared with corresponding genotype in the CD-fed group.

### Iron content was higher in the eWAT from WT and *Hamp1*^*-/-*^ mice, after 24 weeks of HFD-feeding

Protein expression of ferritin in the eWAT was significantly higher in *Hamp1*^*-/-*^ mice than in WT mice, after 12 and 24 weeks of feeding, irrespective of the diet fed (Fig 6A and B). The iron content was also elevated in the eWAT from *Hamp1*^*-/-*^ mice (Fig 6C), which also showed more intense staining with Perls’ stain than that from WT mice, irrespective of the type of diet fed (Supplementary Fig 4A). After 24 weeks of HFD-feeding, ferritin protein levels and the iron content in the eWAT were significantly higher in both genotypes than in the corresponding CD-fed mice (Fig 6A to C). The mRNA and protein levels of TfR1 were similar in WT and *Hamp1*^*-/-*^ mice, irrespective of the type of diet fed (Supplementary Fig 4B and C). The iron content in the eWAT from HFD-fed WT mice was significantly and positively correlated with weights of eWAT, IR and triglyceride content in liver and skeletal muscle (Supplementary Table 3).

**Figure 6:**
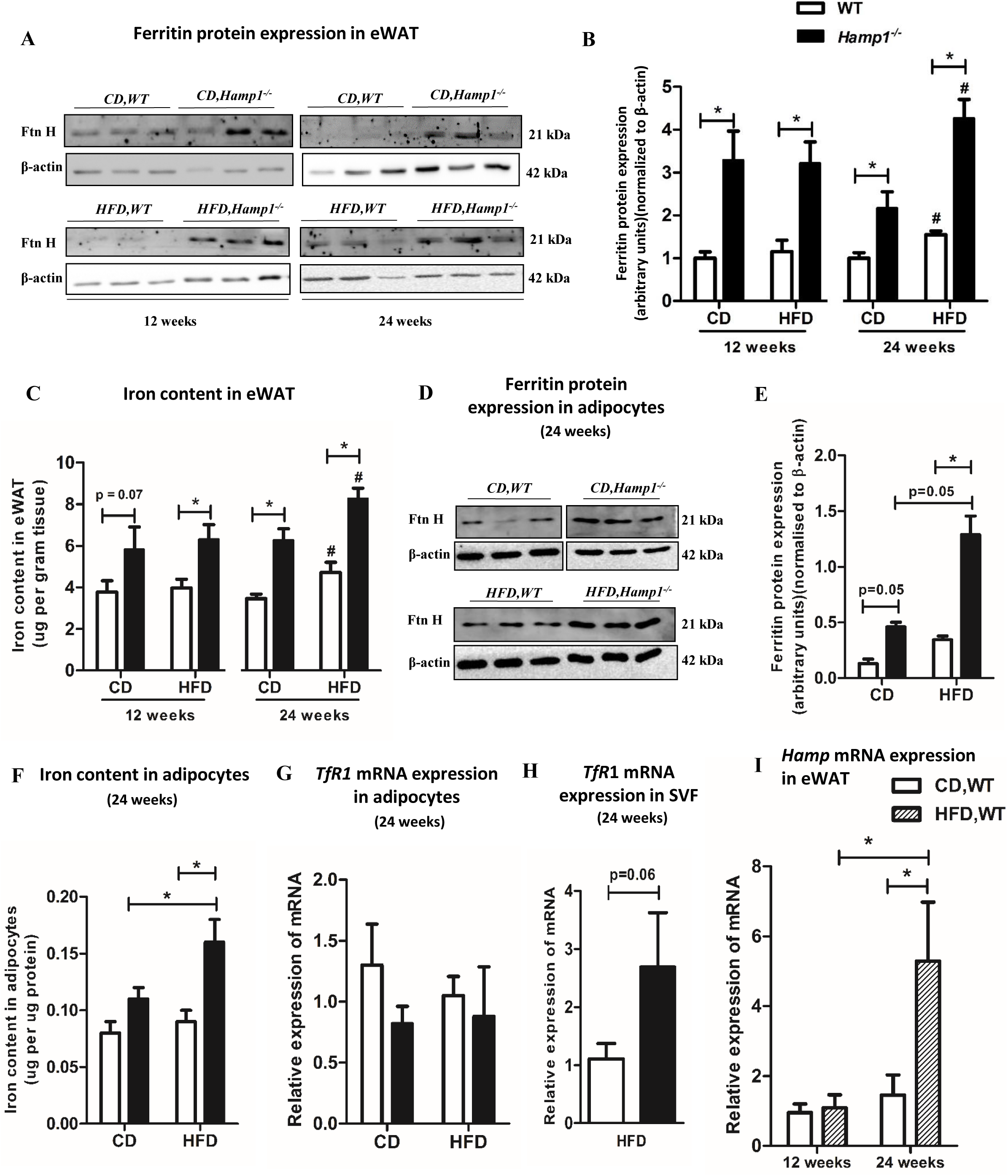
High-fat diet feeding was associated with an increase in iron content in visceral adipose tissue from WT and *Hamp1-/-* mice. Representative images (A), densitometric quantification of western blots for ferritin H (heavy chain) (B) and iron content (C) in the eWAT of wild-type and *Hamp1*^*-/-*^ mice fed CD or HFD for 12 weeks or 24 weeks. Protein levels of ferritin were normalised to that of β-actin, which was used as the loading control. Representative images (D) and densitometric quantification of western blots for ferritin H (E), iron content (F) and mRNA levels of TfR1 (G) in the adipocytes of wild-type and *Hamp1*^*-/-*^ mice fed CD or HFD for 24 weeks. (H) The mRNA levels of TfR1 in the SVF isolated from WT and *Hamp1*^*-/-*^ mice fed HFD for 24 weeks. (I) The mRNA levels of hepcidin in the eWAT of wild-type mice fed CD or HFD for 12 and 24 weeks. Relative expression was calculated by normalising expression of the gene of interest to that of the housekeeping gene, *Hprt*. Results are shown as the mean ± SE, n = 4 to 5 in each group. Data were analysed by Kruskal–Wallis test, followed by Mann–Whitney U test for pair-wise comparisons. \**P* < 0.05 when compared with corresponding WT group; #*P* < 0.05 when compared with corresponding genotype in the CD-fed group.

To determine the distribution of iron within the eWAT, markers of iron status were determined in adipocytes and the SVF isolated from this tissue (from mice fed for 24 weeks). Protein expression of ferritin and iron content in the adipocytes from *Hamp1*^*-/-*^ mice were higher than in those from the WT mice, with significant increases observed in the HFD-fed group (Fig 6D to F). TfR1 mRNA levels (inversely related to its intracellular labile iron pool (34)) were similar in the adipocytes from both genotypes, irrespective of the diet fed (Fig 6G). The mRNA levels of TfR1 in the SVF from HFD-fed *Hamp1*^*-/-*^ mice tended to be higher than in HFD-fed WT mice (p=0.06) (Fig 6H), suggestive of decreased intracellular iron content in the SVF from *Hamp1*^*-/-*^ mice. We attempted to quantify the iron content and ferritin protein content in the SVF to confirm this. However, the quantity of SVF fractions obtained from the eWAT was found to be inadequate for these measurements. Hepcidin (*Hamp1*) gene expression was significantly higher in the eWAT of HFD-fed WT mice than in WT mice fed the CD for 24 weeks or HFD for 12 weeks (Fig 6I).

After 12 and 24 weeks of feeding, the iron content in the liver and skeletal muscle were significantly higher in the *Hamp1*^*-/-*^ mice than the WT mice, irrespective of the type of diet fed (Supplementary Fig 4D and E). In WT mice, HFD feeding for 24 weeks resulted in significantly decreased hepatic iron levels, compared to corresponding CD-fed WT mice. Serum levels of iron tended to be higher in *Hamp1*^*-/-*^ mice than in WT mice at the end of 12 weeks of feeding (either diet) and after CD-feeding for 24 weeks; they were significantly higher in the *Hamp1*^*-/-*^ mice fed the HFD for 24 weeks (Supplementary Fig 4F). The splenic iron content was found to be significantly lower in *Hamp1*^*-/-*^ mice than in WT mice (Supplementary Fig 4G), consistent with an earlier report (25).

## DISCUSSION

C57Bl/6J mice on a HFD have been reported to develop IR by 8 to 12 weeks of feeding. The IR has been shown to increase progressively and become maximal between 20 and 24 weeks (8,26,35). Thus, 12 and 24 weeks (the time points used in the present study) represent periods of early and established HFD-induced IR, respectively.

Increased body iron stores have been reported to be a risk factor for the development of IR (4,36), but its precise role in the development of diabetes is unclear. Ramey et al (37) studied this in hepcidin knock-out mice, which have increased body iron stores. They used 12 months old mice, with a mixed genetic background (129 and C57BL6). At this age, they found that the animals did not spontaneously develop glucose intolerance. They opined that determining whether altered glucose homeostasis would occur on such mice backcrossed with the C57BL6 strain would be of interest, as this strain is considered to be more prone to development of diabetes mellitus (37). The *Hamp1*^-/-^ mice used in the present study had been backcrossed onto the C57Bl/6J strain for 10 generations; using such mice was an attempt to address the lacuna identified by Ramey et al (37). We found that AUCs for the ITT were similar in WT and *Hamp1*^-/-^ mice, when fed CD for 12 and 24 weeks. This observation that *Hamp1*^-/-^ mice did not spontaneously develop IR, despite elevated body iron stores, is in keeping with that by Ramey et al (37) and has shown that the precise background strain of the mice involved did not affect it. However, there was no information (prior to the present study) whether these mice are more susceptible than WT mice to develop IR, in response to HFD-feeding.

In response to 24 weeks of HFD-feeding, *Hamp1*^-/-^ mice developed less IR than HFD-fed WT mice. Since inflammation in eWAT is known to play an important role in the development of HFD-induced IR (8), this aspect was explored. TNF-α and IL-6 are two major pro-inflammatory cytokines, implicated in the initiation and progression of IR (38). In the present study, HFD-feeding resulted in increased expression of TNFα in the eWAT of WT mice; its expression in the eWAT of corresponding *Hamp1*^*-/-*^ mice was significantly lower. However, mRNA (but not protein) levels of IL-6 were higher in the eWAT of HFD-fed *Hamp1*^*-/-*^ mice. IL-6 has been shown to play a critical role in the induction of an anti-inflammatory phenotype in ATMs (39). It is possible that this may contribute to an anti-inflammatory milieu in adipose tissue, as discussed below.

Resident macrophages in adipose tissue have a predominantly anti-inflammatory (M2-like) phenotype (40). In the obese state, circulating monocytes are recruited to adipose tissue, probably mediated by the chemokine, MCP-1 (26,41). These cells infiltrate adipose tissue and differentiate into pro-inflammatory (M1-like) macrophages (11). In the present study, after 24 weeks of HFD-feeding, WT mice had increased numbers of macrophages infiltrating the eWAT, with the macrophages having a marked pro-inflammatory phenotype. On the other hand, eWAT from *Hamp1*^*-/-*^ mice exhibited significantly less MCP-1 expression and lower macrophage infiltration, in response to HFD feeding. Moreover, these macrophages exhibited an anti-inflammatory phenotype. This was associated with a higher number of smaller-sized adipocytes, reduced lipolysis in eWAT and less ectopic lipid accumulation in liver and skeletal muscle in the knock-out mice. These events may account for the lower extent of insulin resistance observed in HFD-fed *Hamp1*^*-/-*^ mice than in WT mice on the same diet. Interestingly, liver triglyceride levels were higher in *Hamp1*^*-/-*^ mice fed CD for 12 and 24 weeks, compared to corresponding WT mice. We suggest this may be on account of enhanced insulin signaling and lipogenesis that has been shown to occur in primary hepatocytes from *Hamp1*^*-/-*^ mice, which we have reported previously (42).

Although iron levels in eWAT were similar in WT and *Hamp1*^*-/-*^ at baseline (0 weeks of feeding) (Supplementary Table 1), its levels were elevated in *Hamp1*^*-/-*^ mice after 12 and 24 weeks, irrespective of the type of diet fed (Fig 6C). However, the iron content in the adipose tissue of mice fed the HFD for 24 weeks was higher than those on the CD for the same duration, irrespective of genotype, suggesting a role for HFD in this effect. Such elevations in iron content may underlie the induction in hepcidin seen in the adipose tissue of WT mice fed the HFD for 24 weeks (Fig 6I). It has been reported that HFD-induced obesity is associated with increased accumulation of iron in ATMs (20). Retention of iron has been reported to contribute to the development of a pro-inflammatory phenotype in macrophages (43,44). In the absence of hepcidin, however, macrophages are likely to be iron-depleted; such an effect is known to occur in macrophages, when hepcidin is deficient (25). It has also been reported that acute iron chelation in human macrophages reduced LPS-induced pro-inflammatory polarization (45). These reports are consistent with our observations of increased TfR1 expression in the SVF fraction (suggestive of decreased intracellular iron content) in *Hamp1*^*-/-*^ mice (Fig 6H) and less inflammation in the adipose tissue (fewer pro-inflammatory M1-like macrophages in the eWAT of the knock-out mice, than in WT mice). They suggest that depletion of iron in macrophages in HFD-fed *Hamp1*^*-/-*^ mice may play a role in decreasing adipose tissue inflammation and subsequent insulin resistance in this setting.

Orr et al (22) studied ATMs isolated from WT C57BL6 mice fed low- and high-fat diets (which had iron contents similar to what was used in the present study). They studied a subset of these ATM, which were natively ferromagnetic, and showed that such cells developed a pro-inflammatory phenotype in response to HFD-feeding. There are several differences in the methodologies employed in their study and the present one. For example, they fed their mice only for 16 weeks, while in the present study the mice were fed for a longer period of 24 weeks. The ferromagnetic ATM they studied were a subset of the total ATMs in the adipose tissue; we did not study such a subset; we studied the total SVF isolated from the adipose tissue. Data on the iron concentrations in the fractions studied have been expressed differently in the 2 studies, precluding direct comparisons of the actual intracellular concentrations involved. However, both studies have suggested that iron content in ATMs may have a role in regulation of insulin sensitivity in the body. Further work would be required to provide more insights into this.

A low macrophage iron content, hepcidin deficiency or altered iron homeostasis have been suggested to account for insulin sensitivity reported in HH mice (18,34,43,46). In the present study, HFD-fed *Hamp1*^*-/-*^ mice showed attenuated inflammation in eWAT and less IR at 24 weeks (at which time point hepcidin was induced in the eWAT of HFD-fed WT mice). We propose that lack of hepcidin in the knockout mice resulted in decreased iron content in the macrophages (due to overexpression of ferroportin that occurs in the absence of hepcidin), predisposing them to become more anti-inflammatory in nature and thereby result in attenuated inflammation in eWAT. Our findings, thus suggest a role for hepcidin in modulating HFD-induced inflammation in the eWAT. Zhang et al (47) have also made such a suggestion recently, based on a different experimental approach. They using shRNA to knock down the hepcidin gene in obese db/db mice. This resulted in decreased inflammation in adipose tissue in these mice. They showed less formation of macrophage extracellular traps (METs) in the adipose tissue in the treated mice and attributed the reduced inflammation to this (47). The results of their work and ours corroborate one another, using different experimental approaches, but propose different mechanisms for the observations.

Although HFD-fed *Hamp1*^-/-^ mice developed less peripheral IR than HFD-fed WT mice, they showed impaired glucose tolerance. A possible reason for this may be an insulin secretory defect in these mice. This is supported by previous reports that have shown insulin secretory defects in *Hfe*^*-/-*^ mice and in patients with HH (17,48). *Hamp1*^-/-^ mice are characterized by severe iron deposition in the exocrine, but not the endocrine, pancreas (49). Although these mice have been reported to develop chronic pancreatitis due to iron toxicity, they showed normal insulin content in pancreas and serum insulin levels (49). Additional work will be required to determine the insulin secretory capacity of beta cells in the pancreas in *Hamp1*^-/-^ mice.

*Hamp1*^-/-^ mice showed less gain in body weight than WT mice in response to HFD, despite similar calorie intake. A similar finding has been reported in *Hjv*^*-/-*^ mice (50). Individuals with hemochromatosis have been reported to have lower body mass index than siblings without hemochromatosis (51). Feeding *Hfe*^*-/-*^ mice with a HFD was found to increase oxygen consumption and heat production in these animals, compared to WT mice (52). It is possible that such a hypermetabolic response may account for the lower gains in body weight in HFD-fed *Hamp1*^-/-^ mice in the present study. Further investigations along these lines would be required to elucidate mechanisms that underlie this.

In conclusion, the present study, which studied the influence of increased body iron stores on the development of IR and inflammation in adipose tissue in response to HFD feeding, has several novel findings to report. Iron-overloaded *Hamp1*^-/-^ mice, used in this study, developed less IR than WT mice, in response to HFD-feeding. This was associated with attenuated inflammation in the eWAT and less hepatic steatosis in these mice. Fewer macrophages infiltrated the eWAT and these exhibited a predominantly M2-like phenotype. We suggest that lowered iron content in macrophages may account for the fewer pro-inflammatory macrophages in the eWAT of the knock-out mice. These findings suggest a novel role for the iron content in ATMs in modulating inflammation in adipose tissue and the possible role of hepcidin in this. Additionally, our data provides early evidence that attenuated inflammation in adipose tissue may account for enhanced insulin sensitivity seen in mice models of, and humans with, hereditary hemochromatosis.

## Data availability statement

The data that support the findings of this study will be available from the corresponding author upon reasonable request.

## Acknowledgements

Centre for Stem Cell Research (CSCR), CMC, Vellore, for use of its flow cytometry core facility.

## Conflict of interest

The authors declare there are no conflict of interests

## Funding

Research grants awarded to MJ by the Science and Engineering Research Board (SERB), Government of India (EMR/2015/000502) and a fluid research grant from Christian Medical College, Vellore (IRB no. 8795, 19^th^ March 2014). JVJ was supported by the SERB grant and then by a Senior Research Fellowship (SRF) from the Indian Council of Medical Research (ICMR), India.

## Contribution statement

JVJ carried out experiments, analyzed the data and wrote the manuscript. JV contributed to conceptualization and design of the study, carried out experiments, analyzed the data and wrote the manuscript. NMJ carried out some of the assays. SV provided the *Hamp1*^-/-^ mice. SV, JCD and ATM critically reviewed results and approved the final version of the manuscript. MJ conceptualized and designed the study, procured funding for the work, analyzed data and wrote the manuscript.

## Notes

### Competing Interest Statement

The authors have declared no competing interest.

