## Supplementary figure for "Insulin resistance and adipose tissue inflammation induced by a high-fat diet are attenuated in the absence of hepcidin"

### Slide 1
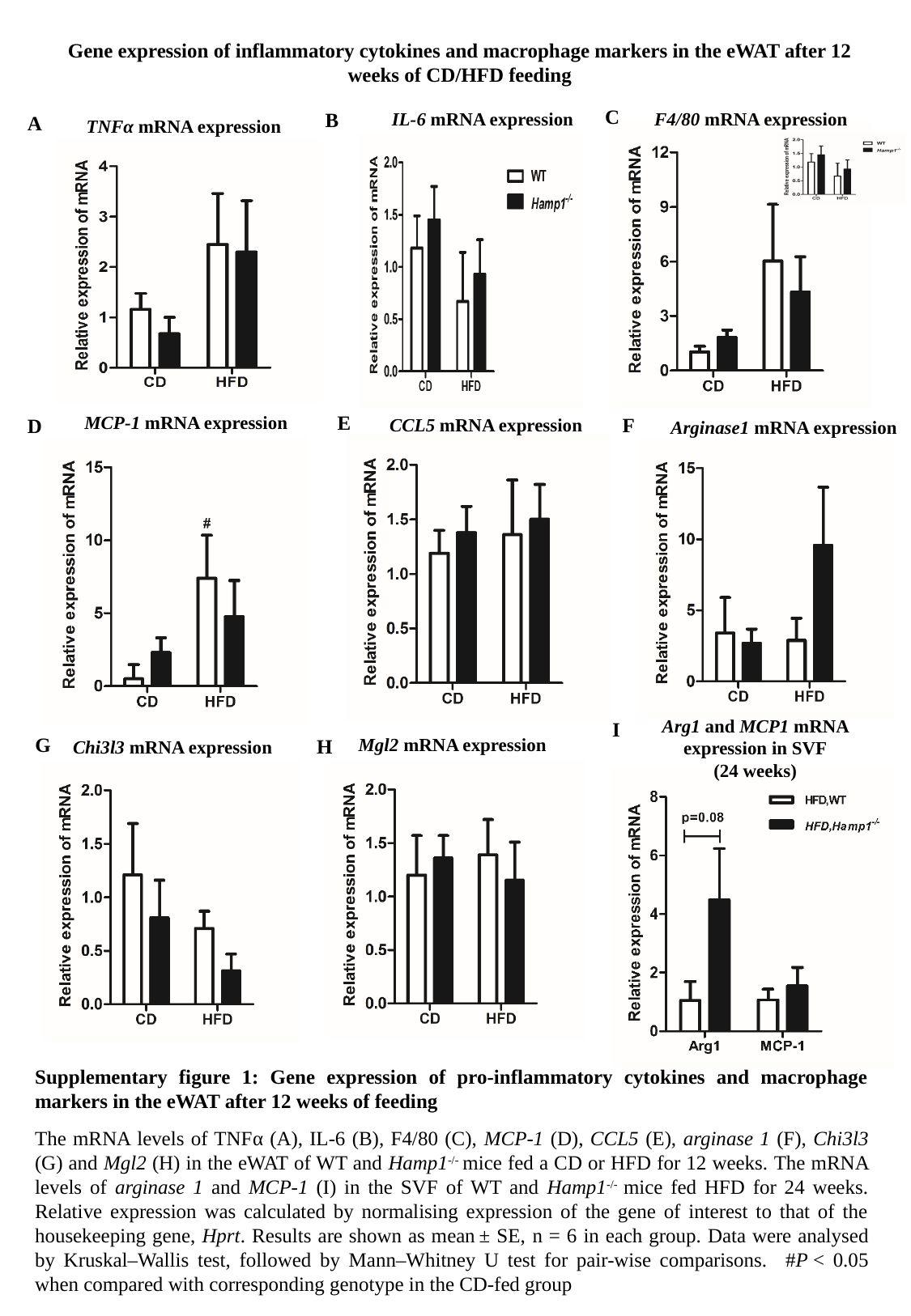

Gene expression of inflammatory cytokines and macrophage markers in the eWAT after 12 weeks of CD/HFD feeding
C
IL-6 mRNA expression
B
F4/80 mRNA expression
A
TNFα mRNA expression
E
MCP-1 mRNA expression
F
 CCL5 mRNA expression
D
Arginase1 mRNA expression
Arg1 and MCP1 mRNA expression in SVF
(24 weeks)
I
G
Mgl2 mRNA expression
H
Chi3l3 mRNA expression
Supplementary figure 1: Gene expression of pro-inflammatory cytokines and macrophage markers in the eWAT after 12 weeks of feeding
The mRNA levels of TNFα (A), IL-6 (B), F4/80 (C), MCP-1 (D), CCL5 (E), arginase 1 (F), Chi3l3 (G) and Mgl2 (H) in the eWAT of WT and Hamp1^-/-^ mice fed a CD or HFD for 12 weeks. The mRNA levels of arginase 1 and MCP-1 (I) in the SVF of WT and Hamp1^-/-^ mice fed HFD for 24 weeks. Relative expression was calculated by normalising expression of the gene of interest to that of the housekeeping gene, Hprt. Results are shown as mean ± SE, n = 6 in each group. Data were analysed by Kruskal–Wallis test, followed by Mann–Whitney U test for pair-wise comparisons. #P < 0.05 when compared with corresponding genotype in the CD-fed group

### Slide 2
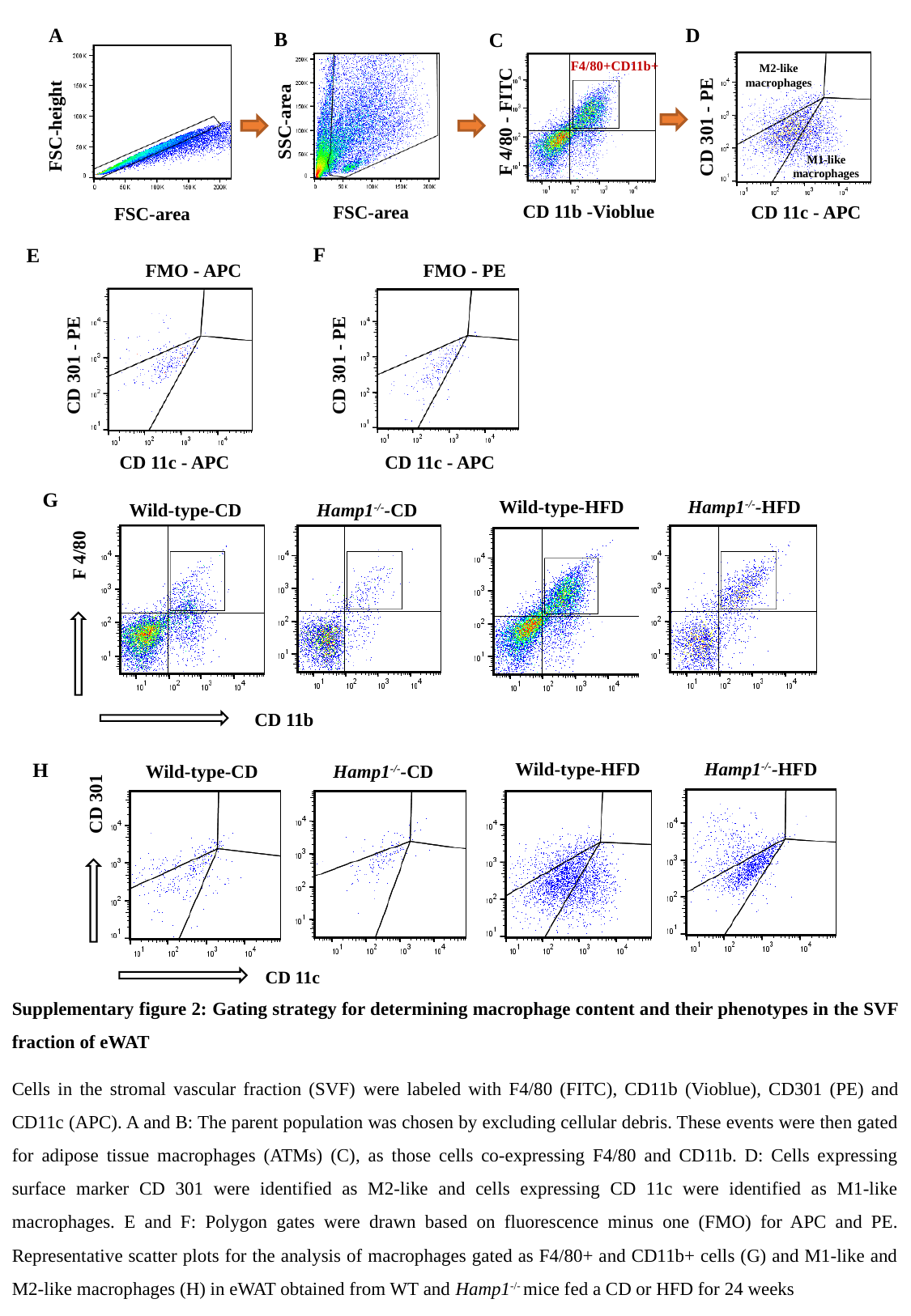

A
D
B
C
F4/80+CD11b+
M2-like macrophages
SSC-area
F 4/80 - FITC
FSC-height
CD 301 - PE
M1-like macrophages
CD 11b -Vioblue
FSC-area
CD 11c - APC
FSC-area
F
E
FMO - APC
FMO - PE
CD 301 - PE
CD 301 - PE
CD 11c - APC
CD 11c - APC
G
Wild-type-HFD
Hamp1^-/-^-HFD
Hamp1^-/-^-CD
Wild-type-CD
F 4/80
CD 11b
H
Wild-type-HFD
Hamp1^-/-^-HFD
Hamp1^-/-^-CD
Wild-type-CD
CD 301
CD 11c
Supplementary figure 2: Gating strategy for determining macrophage content and their phenotypes in the SVF fraction of eWAT
Cells in the stromal vascular fraction (SVF) were labeled with F4/80 (FITC), CD11b (Vioblue), CD301 (PE) and CD11c (APC). A and B: The parent population was chosen by excluding cellular debris. These events were then gated for adipose tissue macrophages (ATMs) (C), as those cells co-expressing F4/80 and CD11b. D: Cells expressing surface marker CD 301 were identified as M2-like and cells expressing CD 11c were identified as M1-like macrophages. E and F: Polygon gates were drawn based on fluorescence minus one (FMO) for APC and PE. Representative scatter plots for the analysis of macrophages gated as F4/80+ and CD11b+ cells (G) and M1-like and M2-like macrophages (H) in eWAT obtained from WT and Hamp1^-/-^ mice fed a CD or HFD for 24 weeks

### Slide 3
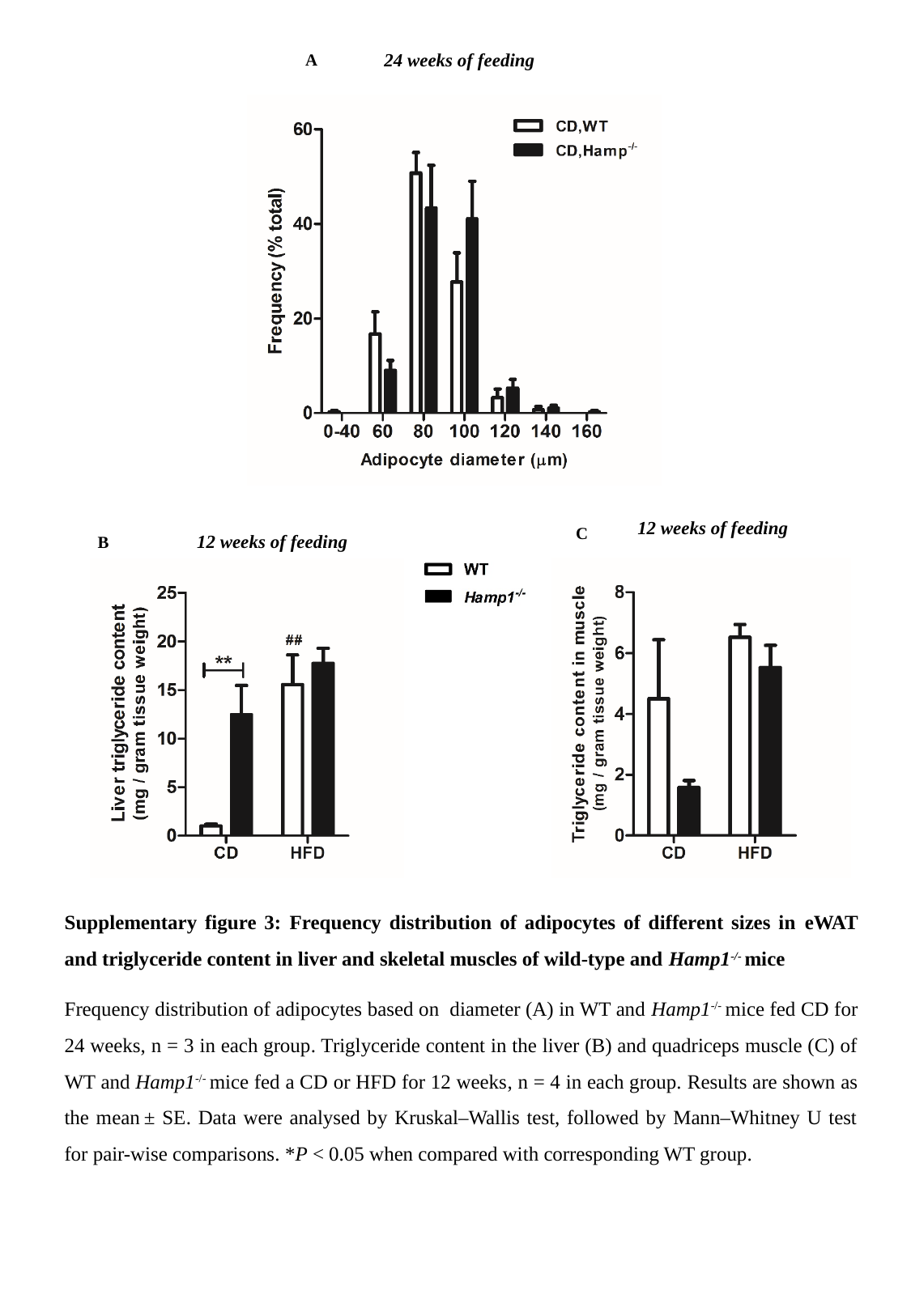

24 weeks of feeding
A
12 weeks of feeding
C
12 weeks of feeding
B
Supplementary figure 3: Frequency distribution of adipocytes of different sizes in eWAT and triglyceride content in liver and skeletal muscles of wild-type and Hamp1^-/-^ mice
Frequency distribution of adipocytes based on diameter (A) in WT and Hamp1^-/-^ mice fed CD for 24 weeks, n = 3 in each group. Triglyceride content in the liver (B) and quadriceps muscle (C) of WT and Hamp1^-/-^ mice fed a CD or HFD for 12 weeks, n = 4 in each group. Results are shown as the mean ± SE. Data were analysed by Kruskal–Wallis test, followed by Mann–Whitney U test for pair-wise comparisons. *P < 0.05 when compared with corresponding WT group.

### Slide 4
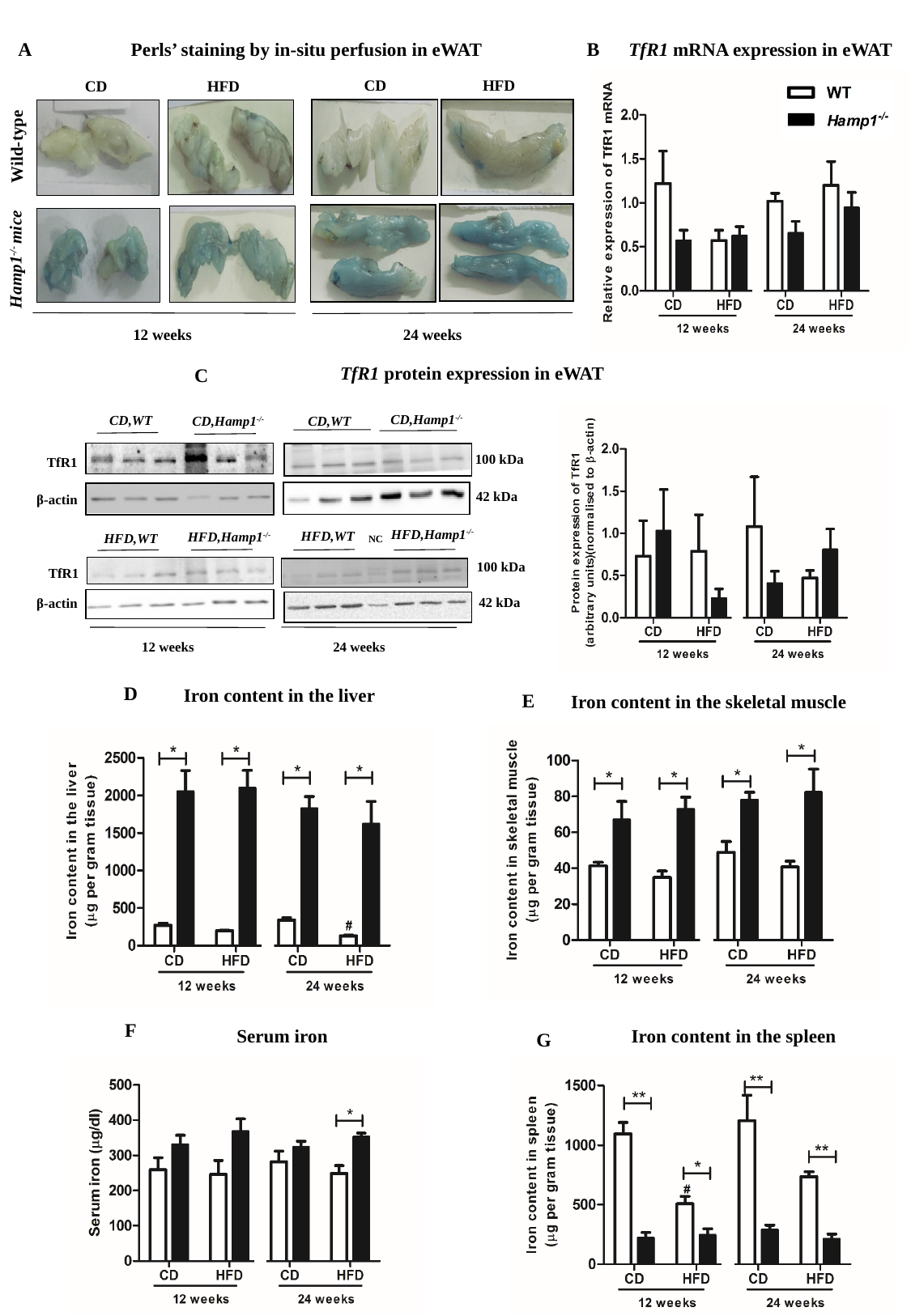

A
B
Perls’ staining by in-situ perfusion in eWAT
TfR1 mRNA expression in eWAT
CD HFD
CD HFD
Wild-type
Hamp1^-/-^ mice
12 weeks
24 weeks
TfR1 protein expression in eWAT
C
CD,Hamp1^-/-^
CD,WT
CD,WT
CD,Hamp1^-/-^
100 kDa
TfR1
42 kDa
β-actin
HFD,Hamp1^-/-^
HFD,WT
HFD,Hamp1^-/-^
HFD,WT
NC
100 kDa
TfR1
42 kDa
β-actin
24 weeks
12 weeks
D
Iron content in the liver
E
Iron content in the skeletal muscle
F
Iron content in the spleen
Serum iron
G

### Slide 5
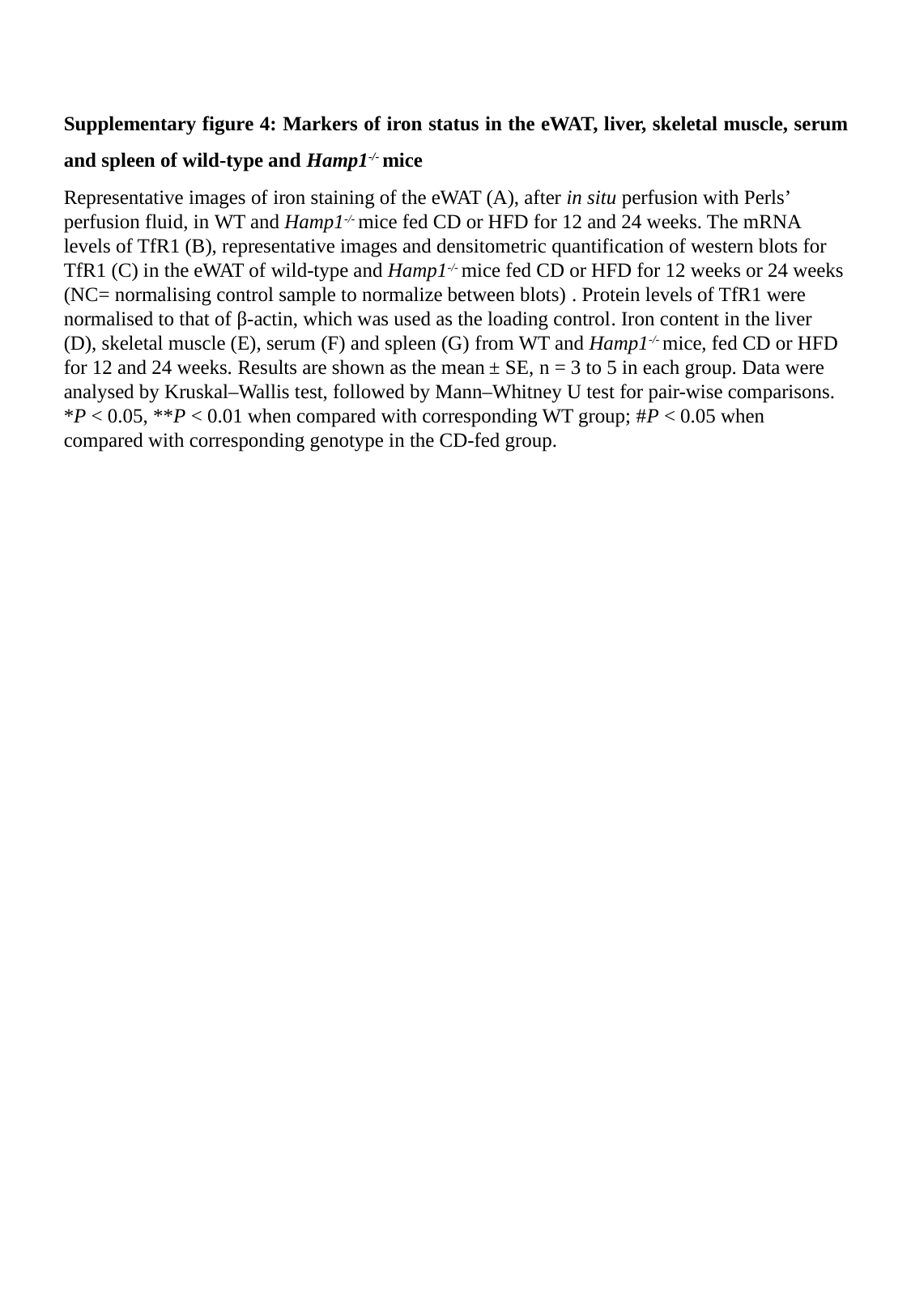

Supplementary figure 4: Markers of iron status in the eWAT, liver, skeletal muscle, serum and spleen of wild-type and Hamp1^-/-^ mice
Representative images of iron staining of the eWAT (A), after in situ perfusion with Perls’ perfusion fluid, in WT and Hamp1^-/-^ mice fed CD or HFD for 12 and 24 weeks. The mRNA levels of TfR1 (B), representative images and densitometric quantification of western blots for TfR1 (C) in the eWAT of wild-type and Hamp1^-/-^ mice fed CD or HFD for 12 weeks or 24 weeks (NC= normalising control sample to normalize between blots) . Protein levels of TfR1 were normalised to that of β-actin, which was used as the loading control. Iron content in the liver (D), skeletal muscle (E), serum (F) and spleen (G) from WT and Hamp1^-/-^ mice, fed CD or HFD for 12 and 24 weeks. Results are shown as the mean ± SE, n = 3 to 5 in each group. Data were analysed by Kruskal–Wallis test, followed by Mann–Whitney U test for pair-wise comparisons. *P < 0.05, **P < 0.01 when compared with corresponding WT group; #P < 0.05 when compared with corresponding genotype in the CD-fed group.
