## Supplementary Table for "Insulin resistance and adipose tissue inflammation induced by a high-fat diet are attenuated in the absence of hepcidin"

**Supplementary table 1: Baseline characteristics of wild-type and *Hamp1^-/-^* mice at 0 weeks of feeding**

| **Variables** | **Wild-type mice** | ***Hamp1^-/-^* mice** | ***P value*** |
| --- | --- | --- | --- |
| **Body weight (g)** | **20.67 ± 0.88** | **21.33 ± 0.33** | **0.63** |
| **eWAT weight (g)** | **0.44 ± 0.05** | **0.31 ± 0.11** | **0.27** |
| **Inguinal SAT weight (g)** | **0.34 ± 0.03** | **0.29 ± 0.07** | **0.31** |
| **Liver weight (g)** | **1.01 ± 0.04** | **1.21 ± 0.11** | **0.12** |
| **Spleen weight (g)** | **0.08 ± 0.01** | **0.09 ± 0.01** | **0.26** |
| **Fasting serum glucose (mg/dl)** | **176.5 ± 37.2** | **210.6 ± 64.2** | **0.82** |
| **Fasting serum insulin (µg/L)** | **0.23 ± 0.10** | **0.21 ± 0.03** | **0.51** |
| **GTT (area under the curve)** | **635.3 ± 94.5** | **506.0 ± 145.8** | **0.02** |
| **ITT (area under the curve)** | **62.2 ± 19.1** | **67.5 ± 25.8** | **0.35** |
| **Liver triglyceride content (mg/gm tissue)** | **10.1 ± 3.16** | **15.89 ± 0.89** | **0.58** |
| **Skeletal muscle triglyceride content (mg/gm tissue)** | **0.63 ± 0.25** | **0.39 ± 0.14** | **0.56** |
| **Serum iron levels (µg/dl)** | **90.9 ± 8.56** | **228.4 ± 22.08** | **0.00** |
| **Ferritin protein expression in liver (arbitrary units)** | **0.43 ± 0.22** | **1.47 ± 0.35** | **0.05** |
| **Ferritin protein expression in skeletal muscle (arbitrary units)** | **0.51 ± 0.24** | **0.63 ± 0.36** | **0.82** |
| **Liver iron content (µg/g)** | **43.6 ± 11.9** | **866.4 ± 171.05** | **0.01** |
| **Epididymal WAT iron (µg/g)** | **4.30 ± 0.41** | **5.68 ± 1.31** | **0.52** |
| **Skeletal muscle iron (µg/g)** | **9.13 ± 0.72** | **12.22 ± 1.40** | **0.12** |
| **Spleen iron (µg/g)** | **439.3 ± 37.9** | **60.2 ± 19.9** | **0.01** |

Data are shown as mean ± SD. Statistically significant correlations are shown as bold. For GTT and ITT, n=20-22 per genotype. For all the other variables, n = 3 to 4 mice per genotype. Data at 0 time is not available for all the parameters studied (for example, estimation of ferritin content in eWAT or in adipocytes or SVF), as the limited quantity of eWAT available in mice of at 8 weeks of age (for both genotypes) did not suffice for all the estimations.

**Supplementary Table 2: Correlational analysis of insulin resistance (as assessed by AUC in ITT) with other parameters of interest**

|  | **WT-CD** | | **WT-HFD** | | ***Hamp1^-/-^* -CD** | | ***Hamp1^-/-^* -HFD** | |
| --- | --- | --- | --- | --- | --- | --- | --- | --- |
|  | r* | *p* | r* | *p* | r* | *p* | r* | *p* |
| *F4/80* mRNA (eWAT) | 0.13 | 0.70 | 0.67 | 0.07 | -0.07 | 0.86 | -0.07 | 0.87 |
| *MCP-1* mRNA (eWAT) | 0.34 | 0.32 | 0.62 | 0.10 | -0.14 | 0.73 | 0.14 | 0.76 |
| *CCL5* mRNA (eWAT) | 0.57 | 0.08 | **0.74** | **0.04** | 0.00 | 1.00 | 0.35 | 0.43 |
| *Arginase1* mRNA (eWAT) | 0.19 | 0.59 | **-0.73** | **0.04** | -0.33 | 0.45 | -0.46 | 0.29 |
| *IL-6* mRNA (eWAT) | -0.35 | 0.31 | **-0.72** | **0.02** | -0.13 | 0.72 | -0.28 | 0.46 |
| Liver triglyceride | -0.08 | 0.81 | **0.84** | **0.002** | 0.45 | 0.23 | 0.26 | 0.48 |
| Liver weight | 0.29 | 0.38 | **0.78** | **0.004** | 0.39 | 0.26 | 0.35 | 0.35 |
| Skeletal muscle triglyceride | -0.09 | 0.79 | 0.61 | 0.07 | 0.45 | 0.24 | 0.11 | 0.76 |

*Spearman’s correlational coefficients are reported. A p value less than 0.05 was taken to indicate statistical significance in all cases. Statistically significant correlations are shown as bold.

**Supplementary Table 3: Correlational analysis of iron content in epididymal white adipose tissue (eWAT) with other parameters of interest**

|  | **WT-CD** | | **WT-HFD** | | ***Hamp1^-/-^* -CD** | | ***Hamp1^-/-^* -HFD** | |
| --- | --- | --- | --- | --- | --- | --- | --- | --- |
|  | r* | *p* | r* | *p* | r* | *p* | r* | *p* |
| Mice body weight | 0.05 | 0.91 | 0.67 | 0.06 | 0.41 | 0.30 | 0.22 | 0.57 |
| eWAT weight | -1.22 | 0.77 | **0.72** | **0.04** | 0.53 | 0.16 | 0.02 | 0.95 |
| *Hamp1* mRNA (eWAT) | -0.63 | 0.17 | 0.52 | 0.28 | - | - | - | - |
| ITT (AUC) | -0.73 | 0.10 | **0.89** | **0.03** | 0.56 | 0.14 | -0.12 | 0.97 |
| *F4/80* mRNA (eWAT) | -0.05 | 0.90 | 0.81 | 0.05 | 0.08 | 0.87 | 0.28 | 0.53 |
| *MCP-1* mRNA (eWAT) | -0.43 | 0.33 | 0.81 | 0.05 | 0.09 | 0.87 | **0.82** | **0.02** |
| Liver triglyceride | 0.08 | 0.84 | **0.94** | **0.001** | 0.37 | 0.40 | 0.15 | 0.71 |
| Skeletal muscle triglyceride | 0.34 | 0.40 | **0.84** | **0.01** | 0.25 | 0.58 | 0.04 | 0.86 |

*Spearman’s correlational coefficients are reported. A p value less than 0.05 was taken to indicate statistical significance. Statistically significant correlations are shown as bold.

Supplementary Table 4: Antibodies used for flow cytometry

| Antigen | Fluorochrome | Source | Species | Dilution | Staining concentration (in µg/ml) |
| --- | --- | --- | --- | --- | --- |
| F4/80 | FITC | Miltenyi Biotech, Germany | Mouse | 1:100 | 0.2ug/ml |
| CD11b | Vioblue |  | Mouse | 1:100 | 0.2ug/ml |
| CD301 | PE |  | Mouse | 1:100 | 0.2ug/ml |
| CD11c | APC |  | Mouse | 1:100 | 0.2ug/ml |

Supplementary Table 5: Primary and secondary antibodies used for western blot

| **No** | **Protein** | **Primary antibody (source and dilution)** | **Primary antibody dilution** | **Secondary antibody**  **(1:5000 dilution)** |
| --- | --- | --- | --- | --- |
| 1 | Ferritin -H | sc-25617; Santa Cruz Biotechnology, Dallas, TX, USA | 1:500 | Anti-rabbit HRP conjugate (Thermo Fischer, Waltham, MA, USA) |
| 2 | TfR1 | #13-6800; Invitrogen, Carlsbad, CA, USA | 1:1000 | Anti-mouse HRP conjugate (Thermo Fischer, Waltham, MA, USA) |
| 3 | Phospho-Akt  (Ser 473) | #9271; Cell signaling technology, Danvers, MA, USA | 1:1000 | Anti-rabbit HRP conjugate (Thermo Fischer, Waltham, MA, USA) |
| 4 | Total Akt | #9272; Cell signaling technology, Danvers, MA, USA | 1:1000 | Anti-rabbit HRP conjugate (Thermo Fischer, Waltham, MA, USA) |
| 15 | Beta actin | #A5316; Sigma-Aldrich, St. Louis, MO, USA | 1: 5000 | Anti-mouse HRP conjugate (Thermo Fischer, Waltham, MA, USA) |

Supplementary Table 6: Primers used for quantitative PCR

| **Sl.**  **No.** | **Gene** | **Primer sequence (mouse)** | **Amplicon size (bp)** |
| --- | --- | --- | --- |
| 1 | *Tfr-1* | 5’-GAG GCG CTT CCT AGT ACT CC-3’  3’-CTT GCC GAG CAA GGC TAA AC-5’ | 121 |
| 2 | *Arginase-1* | 5’-CTC CAA GCC AAA GTC CTT AGA G-3’  5’-AGG AGC TGT CAT TAG GGA CAT C-3’ | 185 |
| 3 | *Chi3l3* | 5’-AGA AGG GAG TTT CAA ACC TGG T-3’  5’-GTC TTG CTC ATG TGT GTA AGT GA-3’ | 109 |
| 4 | *TNFα* | 5’-AAG CCT GTA GCC CAC GTC GTA-3’  5’-GGC ACC ACT AGT TGG TTG TCT TTG-3’ | 122 |
| 5 | *F4/80* | 5’-CTT TGG CTA TGG GCT TCC AGT C-3’  5’-GCA AGG AGG ACA GAG TTT ATC GTG-3’ | 165 |
| 6 | *CCL5* | 5’-TGC CCT CAC CAT CAT CCT CAC T-3’  5’-GGC GGT TCC TTC GAG TGA CA-3’ | 194 |
| 7 | *IL-6* | 5’-CCA CTT CAC AAG TCG GAG GCT TA-3’  5’-GCA AGT GCA TCA TCG TTG TTC ATA C-3’ | 112 |
| 8 | *Mgl2* | 5’-TTA GCC AAT GTG CTT AGC TGG-3’  5’-GGC CTC CAA TTC TTG AAA CCT-3’ | 102 |
| 9 | *MCP-1* | 5’-AGG TCC CTG TCA TGC TTC TGG-3’  5’-CTG CTG CTG GTG ATC CTC TTG-3’ | 169 |
| 10 | *Hprt* | 5’- CTGGTTAAGCAGTACAGCCCCAA-3’  5’- CGAGAGGTCCTTTTCACCAGC-3’ | 65 |

Supplementary Table 7: quantitative PCR validation data

| **Sl. No** | **Gene** | **Standard curve slope** | **R^2^ of standard curve** | **Linear dynamic range (cDNA**  **dilution)** | **Standard deviation of Ct at lower limit of dynamic range** | **Primer dimer (melting curve analysis)** | **Ct of amplification (if any) in the NTC** |
| --- | --- | --- | --- | --- | --- | --- | --- |
| 1 | *Tfr-1* | -3.689 | 0.999 | 1:5 to 1:625 | 0.06 | None | - |
| 2 | *Arginase-1* | -3.11 | 0.994 | 1:5 to 1:625 | 0.02 | None | - |
| 3 | *Chi3l3* | -3.04 | 0.998 | 1:5 to 1:3125 | 0.06 | None | - |
| 4 | *TNFα* | -3.236 | 0.999 | 1:5 to 1:3125 | 0.07 | None | - |
| 5 | *F4/80* | -3.325 | 0.999 | 1:5 to 1:3125 | 0.08 | None | - |
| 6 | *CCL5* | -3.384 | 0.998 | 1:5 to 1:3125 | 0.09 | None | 33 |
| 7 | *IL-6* | -3.572 | 0.969 | 1:5 to 1:3125 | 0.07 | In NTC | 35 |
| 8 | *Mgl2* | -3.188 | 0.996 | 1:5 to 1:3125 | 0.13 | In NTC | - |
| 9 | *MCP-1* | -2.985 | 0.999 | 1:5 to 1:3125 | 0.02 | None |  |
| 10 | *Hprt* | -3.409 | 0.998 | 1:5 to 1:3125 | 0.02 | None | - |
